# Atypical Audio-Visual Neural Synchrony and Speech Processing in children with Autism Spectrum Disorder

**DOI:** 10.1101/2024.04.19.590044

**Authors:** Xiaoyue Wang, Sophie Bouton, Nada Kojovic, Anne-Lise Giraud, Marie Schaer

## Abstract

**Background:** Children with Autism Spectrum Disorders (ASD) often exhibit communication difficulties that may stem from basic auditory temporal integration impairment but also be aggravated by an audio-visual integration deficit, resulting in a lack of interest in face-to-face communication. This study addresses whether speech processing anomalies in young (mean age 3.09-year-old) children with ASD are associated with alterations of audio-visual temporal integration.

**Methods:** We used high-density electroencephalography (HD-EEG) and eye tracking to record brain activity and gaze patterns in 31 children (6 females) with ASD and 33 typically developing (TD) children (11 females), while they watched cartoon videos. Neural responses to temporal audio-visual stimuli were analyzed using Temporal Response Functions model and phase analyses for audiovisual temporal coordination.

**Results:** The reconstructability of speech signals from auditory responses was reduced in children with ASD compared to controls, but despite more restricted gaze patterns in ASD it was similar for visual responses in both groups. Speech reception was most strongly affected when visual speech information was also present, an interference that was not seen in TD children. These differences were associated with a broader phase angle distribution (exceeding pi/2) in the EEG theta range in autistic children, signaling reduced reliability of audio-visual temporal alignment.

**Conclusion:** These findings show that speech processing anomalies in ASD do not stand alone and that they are associated already at a very early development stage with audio-visual imbalance with lousier auditory response encoding and disrupted audio-visual temporal coordination.

## Introduction

Newborns are immediately attracted to the human voice. In utero exposure to speech sounds enables them to accurately discriminate speech sounds at birth (1–3). Since vision develops with a delay relative to hearing, babies only progressively discover that vocal stimuli are related to facial movements. Unlike typically developing (TD) children, children with Autism Spectrum Disorders (ASD) do not show this primary interest in speech (4–7). Instead, they tend to engage in slow and repetitive visual exploration of their environment, which has been suggested to lead to atypical interests over time (8–13). Focusing on visual aspects of their surroundings allows children with ASD to explore the world at their own pace, keeping them away from highly dynamic stimuli such as speech and biological motion (14–16), which are often perceived as overwhelming (17,18).

A basic auditory dysfunction in ASD might lead to speech-processing anomalies that in turn cascade into a decreased interest in speech (19–24). Atypical speech processing becomes apparent very early in development, and the fact that early neural anomalies, such as delta, theta, and gamma oscillations, accurately predict the severity of future language deficits could suggest that they are causal to later difficulties in language comprehension and production (25–28). The tendency of children with ASD to prefer static or slow visual processing (29–32) possibly exacerbates speech reception challenges by counteracting dynamic audio-visual interaction, a crucial process for speech reception in ecological (e.g., noisy) environments (33–36). Accordingly, exceedingly long integration time windows for audio and visual stimuli have been reported in children with ASD (31,32), implying disturbed integration of audio and visual stimuli in ASD.

Two essential mechanisms participate in audio-visual integration. The first one is the relative timing of auditory and visual stimuli: when falling within approximately 250 ms of each other, they are often perceived as a single event, potentially influencing each other (e.g. the McGurk effect (37,38)). The second mechanism is the resynchronization provoked by the stimulus in one sensory modality affecting neural responses in the other one (39–44). Orofacial visual movements typically precede the onset of speech, leading to a resynchronization that sharpens the auditory speech response (39,45). And independent from the integration/fusion of the exact visual and speech content, visual resynchronization enhances speech processing by boosting the tracking of the speech’s syllabic structure. While audio-visual temporal integration anomalies in ASD are well documented (31,32,46–48), audio-visual dynamic synchronization anomalies remain hypothetical.

Auditory and visual sensory processing both operate rhythmically (49–53). Visual speech information (lip movements) is characterized by a dominant 2-7 Hz rhythm (theta band (54)) and these quasiperiodic visual cues influence speech perception by modulating auditory neuronal oscillations within the same theta range at about 5 Hz (53–58), corresponding to the typical audio-visual (AV) integration temporal time around 250ms (39,61–64). The reset of auditory neural oscillations triggered by visual input (65) rhythmically enhances auditory processing (66), a phenomenon that is already observable in typical children (67). Despite the documented presence of auditory processing anomalies in ASD around 3-year-old (24), we still ignore whether they are associated with dynamic audio-visual synchrony anomalies.

This study fills this gap by investigating with high-density EEG the dynamics of auditory and visual processing in young children with and without ASD, aged 1.13 to 5.56 years old, under naturalistic audio-visual conditions, i.e. when children are watching a popular cartoon adapted to their age. The goal is to compare the quality of the neural encoding/decoding of dynamic auditory and visual stimuli and audio-visual temporal coordination across groups.

## Methods

### 1 Participants

Participants were selected from the Geneva Autism Cohort, a longitudinal study that aims at better understanding the developmental trajectories in young children with ASD. This cohort’s protocol has been detailed in previous studies (22,62,63). In this study, we used clinical and behavioral assessments, as well as the electroencephalogram (EEG) recorded simultaneously with eye-tracking when children were watching popular cartoon videos.

The sample comprised 31 children diagnosed with ASD (6 females, mean age = 3.09 years, SD = 0.91, age range: 1.74 - 5.14) and 32 TD peers (11 females, mean age = 2.95 years, SD = 1.31, age range: 1.31 - 5.56). Selection criteria for all participants included: age below 6 years, data collected during the participant’s initial visit (i.e. at autism diagnosis for the autistic group), clear and accurate markers associated with movie onset, usable raw data for four different movies, and focus on the screen throughout all recordings. The age difference between the two groups was not significant (Kolmogorov-Smirnov D = 0.28, p = 0.17).

ASD clinical diagnoses were meticulously confirmed using standardized tools: either the Autism Diagnosis Observation Schedule-Generic (ADOS-G) (70) or the Autism Diagnosis Observation Schedule, Second Edition (ADOS-2) (71). Recruitment of participants occurred through specialized clinical centers and community-wide announcements. For the TD group, exclusion criteria included any suspicion of atypical psychomotor development, a history of neurological or psychological disorder, or having a first-degree relative with an autism diagnosis.

Informed consent was obtained from the parents of all participants prior to inclusion in the study. The research was conducted with the ethical standards set forth by the Ethics Committee of the Faculty of Medicine at the University of Geneva Hospital and adhered to the principles outlined in the Declaration of Helsinki.

### 2 Stimuli and Procedure

To explore cortical processing of audio-visual stimuli, we employed a passive and naturalistic task suitable for young children. This task involved viewing an age-appropriate French cartoon “TROTRO” (72–75) (example: https://www.youtube.com/watch?v=jT9C9WCIQr8&t=81s). The selection of “TROTRO” was based on its cognitive accessibility and appeal to the target age group. Importantly, TROTRO, the main character, speaks and interacts verbally with other characters, which allows us to isolate the speech soundtrack and associated visual motion and to probe related brain responses. Participants watched four Trotro episodes, each lasting approximately 2.5 minutes. The videos were presented in a consistent, predetermined order to all participants. To monitor the participant’s visual engagement with the stimulus, Tobii Studio (Tobii® Technology, Sweden) was used. The screen for the video display was configured with dimensions of 1200 pixels in height (29°38’ visual angle) and 1920 pixels in width (45°53’), with a refresh rate of 60 Hz. This setup was optimized for clear and comfortable viewing in children. Participants were seated at an optimal distance of approximately 60 cm from the screen (Figure 1A).

**Figure 1.**
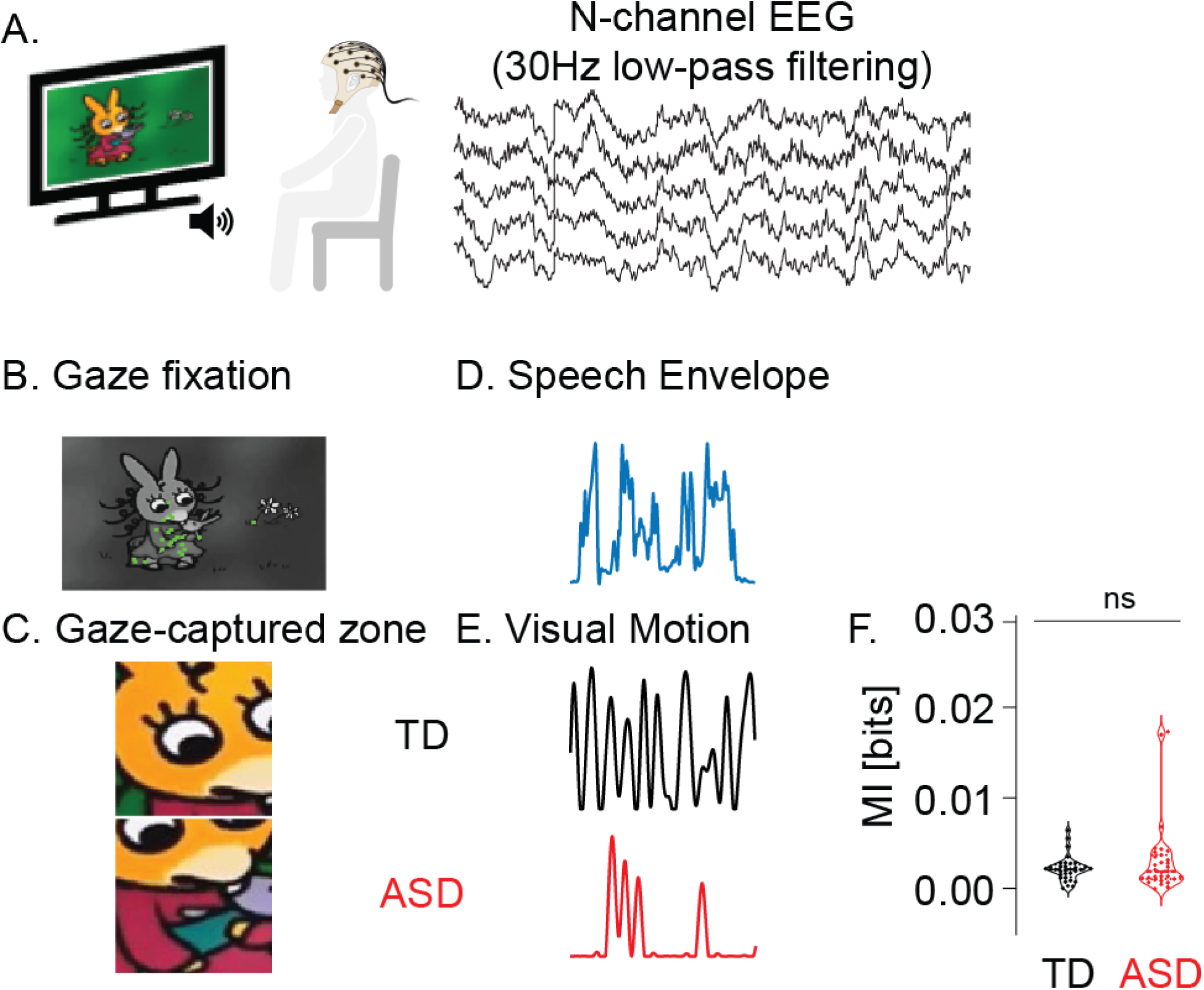
Overview of Experimental Procedures and Features of Interest. (A) Experimental procedures (B) Gaze fixation. Example of individual gaze fixation points (green dots) on a black and white image; (C) Example of gaze-captured screen areas. Depiction of screen areas captured by the gaze of participants in ASD (Autism Spectrum Disorder) and TD (Typically Developing) groups. (D) Example of a stimulus speech envelope from the cartoon soundtrack. (E) Visual motion corresponds to the same stimulus in each group. (F) Comparison of speech envelope and visual motion. Mutual Information (MI) between ASD and TD groups (ns. p>0.05).

#### 2.1 Eye-tracking acquisition and analysis

Gaze data were collected using the Tobii TX300 eye-tracking system (https://www.tobiipro.com), which operates at a sampling rate of 300 Hz. This high-frequency data collection was instrumental in assessing participants’ visual exploration patterns during the cartoon viewing. The cartoon was displayed in a frame that provided a visual angle of 26°47’(height) × 45°53’(width). Calibration was performed using a child-friendly procedure integrated into the Tobii system, specifically designed to engage young participants. This calibration was critical for accurate gaze position tracking and was repeated as necessary, particularly in instances where the eye-tracking device showed any discrepancies in detecting the participant’s gaze. To maintain consistency and reliability in data quality, we ensure constant lighting conditions in the testing room throughout all sessions. Special consideration was given to the youngest participants, who were seated on their parent’s lap when they felt more comfortable in this setting, a strategy that effectively minimized potential head and body movements that could interfere with accurate data collection. For data analysis, we employed the Tobii IV-T Fixation filter (13,76). This tool is specifically designed to extract fixation data, providing us with precise and reliable measures of visual attention and engagement.

#### 2.2 Audio and visual stimuli

We edited the original movie soundtrack using Audacity v.2.2.1 (Audacity Team, 2021) to isolate speech excerpts while removing extraneous background noise such as birds’ singing and musical interludes.

Having extracted the video speech-track, we explored corresponding visual dynamics. Within the video excerpts that contained speech, we distinguished two key components: speech envelope (Figure 1B1) and visual motion (Figure 1B2). To extract the speech envelope, we used the absolute value of the analytic signal (77). The obtained speech envelope was then down-sampled to 1000 Hz and filtered using a zero-phase, fourth-order Butterworth filter with a 40 Hz cutoff. The visual motion was restricted to the participant’s gaze-attended zone. We considered individual gaze-fixation positions and the size of retinal stimuli around the gaze-fixation point. Our analysis took into account the decline in visual acuity and the crowding effects of parafoveal vision (2–5 degrees from the fixation point) (78). The region for capturing visual stimuli through eye gaze was defined as a square with sides the length of an 8-degree diameter centered on the fixation point, aligning with findings on the effective visual span guiding saccades (79). This corresponded to approximately 318 × 318 pixels (Figure 1 A1 & A2). The gaze-capture zone was converted to greyscale, and luminance differences between successive frames were computed following established methodologies (80). The extraction of corresponding visual stimuli involved converting the region of interest of each frame to a grayscale and computing the luminance difference between successive frames. Pixels with a luminance change greater than 10 (a threshold chosen to mitigate video recording noise) were selected, and the average luminance change constructed the visual motion component.

**Figure 2.**
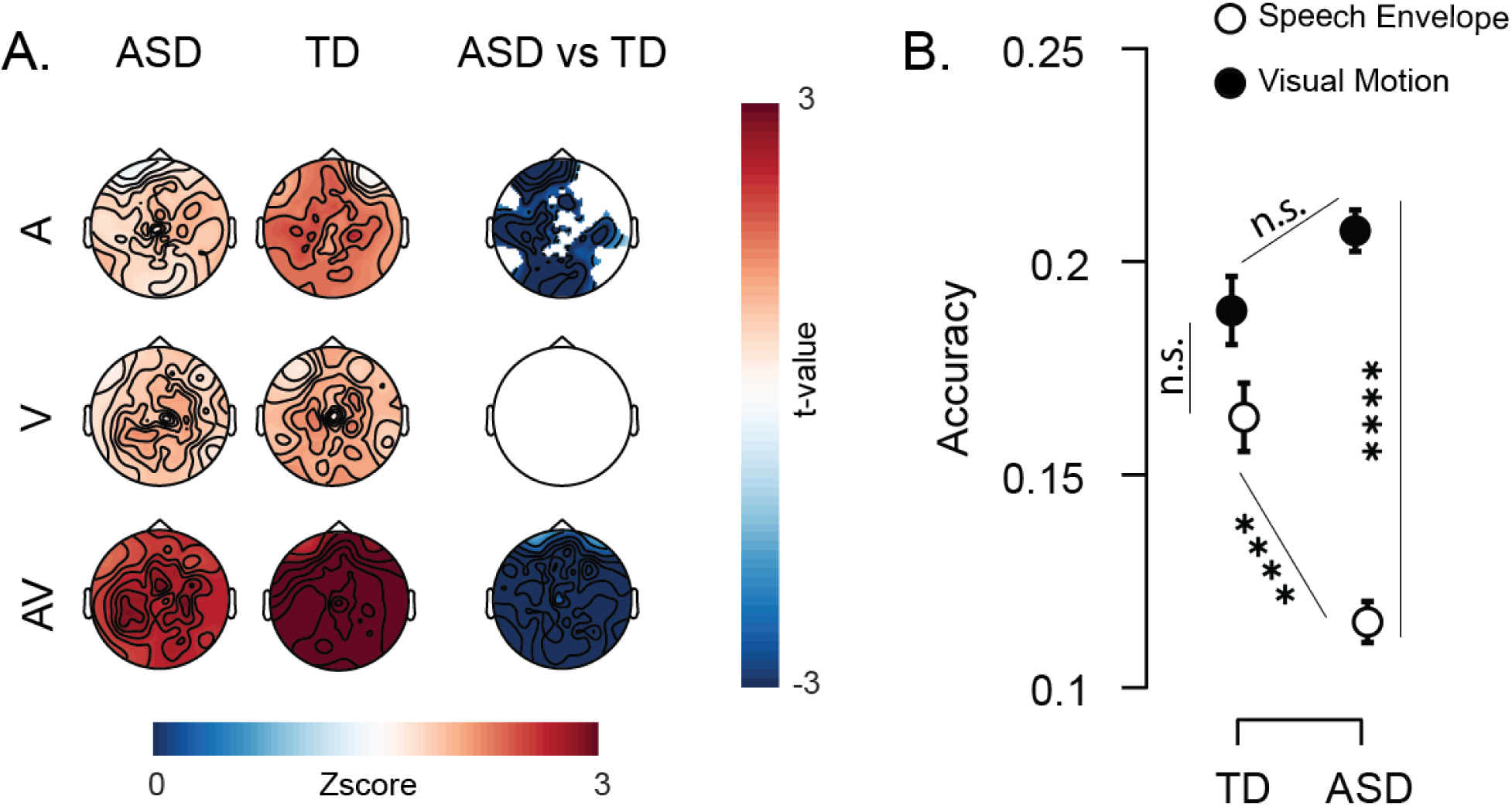
Comparison of audio, visual, and AV models. **(A.) Neural representations** in ASD (left) and TD (middle) groups, for each model (A-speech envelope, V-visual motion, and AV joint) across all scalp electrodes. The right column shows EEG channels where significant group differences are observed using cluster-based nonparametric statistics (p < 0.05; with a positive t-value indicating greater predictability in the ASD group compared to the TD group). **(B.) Stimulus reconstruction accuracy** for speech envelope and visual motion in both groups. Error bars indicate the standard error of the mean. Significance levels are indicated as follows: ‘ns’ for p>0.05 (not significant), * for p <0.05, ** for p<0.01, *** for p<0.001, and **** for p<0.0001.

Visual motion data was upsampled to match the 1000 Hz EEG sampling frequency. Speech envelopes and visual motion corresponding to the same speech-track were aligned and prepared for further analysis. With this method, visual motion is inherently individual-specific, as it relies on the visual exploration of each participant.

#### 2.3 Stimulus features analysis

In order to assess shared information between speech envelope and visual motion, in ASD and TD groups, we calculated mutual information (MI) scores, a dynamic metric, expressed in bits, which quantifies the reduction in uncertainty of one variable when another is observed (81,82). We calculated MI using the *quickMI* function from the Neuroscience Information Theory Toolbox (83). The parameters for this calculation were set to 4 bins, no delay, and a p-value threshold of 0.001 (83). For generating the MI scores, we concatenated all kept excerpts in the same sequence across subjects, separately for each stimulus feature and each group (ASD and TD). This process was followed by a comparative analysis of MI values between groups.

We only included stimuli corresponding to time periods with usable EEG signals. This resulted in slight variations in the stimulus duration for the ASD and TD groups, for which we controlled. No significant disparities in MI scores emerged between the two groups (*t*(1, 61)= 1.250, p = 0.216, Cohen’s d = 0.315, Figure 1C), indicating that these minor differences did not lead to notable group differences in the shared information between the speech envelope and visual motion.

### 3 EEG acquisition and pre-processing

The EEG data were acquired through a 129-electrode (Hydrocel Geodesic Sensor Net (HCGSN) system (Electrical Geodesics, USA) at a sampling rate of 1000 Hz. During recording, the signals were subjected to real-time 0-100 Hz band-pass filtering. The reference electrode was positioned at the vertex (Cz). Data pre-processing was conducted using the EEGLAB v2019 toolbox within the MATLAB environment (84) and Cartool (https://sites.google.com/site/cartoolcommunity/). One hundred and ten channels were kept excluding the cheek and neck to prevent contamination by muscle artifacts. EEG signals were filtered using a zero-phase fourth-order Butterworth bandpass (0.1-70 Hz) and a 50 Hz notch filter to eliminate power line noise. EEG data were visually inspected to remove movement artifact-contaminated periods. Bad channels were first identified and excluded for excessive signal amplitude. Eye blinks, saccades, electrical noise, and heartbeat artifacts were excluded using independent component analysis (ICA). A spherical spline interpolation was used to interpolate the channels contaminated by noise using the ICA-corrected data. Finally, a common average reference was recalculated on the cleaned data, with an additional step of applying a 30Hz low-pass filtering (80). To ensure that all the EEG signals and stimulus features were on a similar scale and thus comparable, we normalized both the EEG signals and stimulus features (i.e. speech envelope and visual motion) using the *nt_normcol* function (Noisetools: http://audition.ens.fr/adc/NoiseTools/).

### 4 Temporal Response Functions (TRF)

To quantify how well EEG in ASD and TD children linearly varied with the stimulus features, we performed regularized regression (with ridge parameter λ) as implemented in the mTRF toolbox (85). The Temporal Response Function (TRF) captures how the brain’s EEG activity correlates with and responds to changes in the stimulus over time, providing a dynamic mapping of the neural processing of the stimulus features. More precisely, the TRF accounts for the fact that the brain’s response to a stimulus occurs with a certain delay. Specifically, the TRF analysis models the relationship between each stimulus feature (i.e. speech envelope or visual motion) and the brain’s response, particularly the time-lagged aspects of this relationship. The TRF includes a coefficient for each time lag that quantifies the strength and direction (positive or negative) of the brain’s response to the stimulus at that specific time delay. The lags in a TRF are represented as a series of time-points or intervals, typically in milliseconds. For the TRF estimation, we downsampled all signals to a rate of 100 Hz to speed computation.

#### 4.1. Estimation of TRF using forward encoding models

Specifically, we used a forward encoding model approach. Since changes in the EEG signal are expected with an unknown time lag after the stimulus, predictions were computed over a range of time lags between 300 ms earlier than the stimulus and 300 ms later than the stimulus. To single out brain signals involved in auditory and visual dynamics, we constructed two distinct univariate encoding models using the speech envelope and visual motion as independent regressors respectively (labeled A-only and V-only). Further advancing our investigation, we developed a multivariate encoding model, labeled ‘AV-joint’, which integrates both the speech envelope and visual motion as regressors. The integration within this model is operationalized through the assignment of trade-off weights to the AV regressor, which are calibrated to reflect the respective contributions of auditory and visual stimuli. These trade-off weights ensure a balanced representation within the model, enabling for an accurate representation of the brain’s concurrent processing of both modalities. This balanced approach aims to embody effective multisensory integration, preventing the overshadowing of one sensory modality by another.

The comparative analysis of the AV-joint model against the A-only and V-only models is essential to elucidate the incremental benefits of simultaneous audio-visual integration over unimodal processing. By evaluating the performance and predictive accuracy of the multivariate model relative to its univariate counterparts (one focused only on auditory processing and the other on visual processing), we aim to substantiate the hypothesis that the synergistic consideration of auditory and visual stimuli offers a more comprehensive understanding of sensory processing in naturalistic listening environments.

For each group, we created “generic” models that predict the EEG data of an individual participant (nth participant) using a TRF derived from the EEG data of the other participants (the remaining n-1 participants). We thus implemented an *n*-fold leave-one-out cross-validation strategy and optimized the model through a parameter search for the regularization parameter λ.

#### 4.2 Selection of optimal regularization parameter

To mitigate the risk of data overfitting in forward encoding models, we integrated an optimized regularization parameter λ, determined for each stimulus feature. To do so, we trained multiple model iterations on subsets of data comprising n-1 participants. During these iterations, λ was systematically adjusted within a predefined range from 10^0^ to 10^5^, with increments in the exponent of 0.5. The criterion for selecting the optimal λ was maximal predictive accuracy. This was quantitatively assessed by calculating Pearson’s correlation coefficient (r) between the predicted EEG signals and the observed signals, for each electrode and each participant. Once optimal λ was identified, the refined model was used to predict the EEG responses of the nth participant, using an n-fold leave-one-out cross-validation paradigm. This methodology was applied consistently across both ASD and TD groups, ensuring the robustness and reliability of the predictive models within each group.

#### 4.3. Pearson’s correlation coefficient (r)

Pearson’s correlation coefficient (r) was computed to quantify the model’s prediction accuracy. For each electrode, the correlation between the EEG signals and the TRF-predicted signals was calculated across all time points. This process was repeated for each participant for the validation set. The correlation coefficients were then averaged across participants to obtain a mean Pearson’s r-value per electrode. This allowed us to create a scalp-wide map of the model’s prediction accuracy, highlighting areas with the strongest correlation between the predicted and observed EEG responses. The accuracy of EEG predictions derived from the optimized model was then converted into Z-scores for further statistical group comparisons. The Z-score computation involved subtracting the mean of surrogate data and dividing it by the standard deviation of surrogate data. Surrogate distributions were generated by randomly shifting (50 times) the orders of the testing EEG segments, preserving their original temporal structure. In summary, this analysis indicated the accuracy with which stimulus features are predicted from the EEG data for each participant.

To quantitatively compare the accuracy between the ASD and TD groups, we used a cluster-based permutation test with 1000 randomization iterations, following the approach of Maris and Oostenveld (86). Clusters were defined by considering both time and spatial electrode configurations, requiring each cluster to include at least two adjacent electrodes. A pivotal aspect of this approach was ensuring that the cluster-level type-I-error probability remained below the 0.05 threshold. This strategy was effective in controlling the family-wise error rate, maintaining it within the 5% type-I-error rate boundary.

#### 4.4. Stimulus reconstruction using decoding models

We trained decoders using EEG data across a wide temporal range, from −300 to 300 ms relative to the stimulus, aiming for optimal stimulus reconstruction. This analysis involved an iterative process of leave-one-out cross-validation and optimization techniques to refine our decoding accuracy, assessing how effectively the EEG signals could predict the stimulus features (i.e., visual motion and speech envelope) for each group.

To enhance our understanding of the temporal alignment between EEG signals and the stimuli, our methodology evolved to focus on discrete, predefined time-lag intervals for decoder training, rather than a continuous range. By pinpointing discrete intervals and evaluating their decoding success, we identified the most informative time-lag segment that yielded the highest fidelity in stimulus reconstruction. Determining this optimal time-lag interval informs us about the specific moments when the EEG data are most synchronized with the stimulus features, thus shedding light on the precise neural timing critical for effective sensory processing and integration.

For the decoding models, we determined the accuracy of stimulus reconstruction accuracy and the identity of the best time lag using the Kruskal-Wallis test with Dunn’s multiple comparisons test (87). This non-parametric statistical method was chosen for its capability to handle variations in group means and variances across different conditions.

### 5. Low-frequency tracking of audio-visual signal

To explore whether the combined processing of auditory and visual stimuli can be explained by the tracking of audio-visual signals by low-frequency brain activity, we used a coherence-based and phase-based analytical framework (88). This approach probed the interplay between neural responses and stimulus features by comparing their magnitude spectra and the phase relationship. Our focus centered on the delta and theta frequency bands, which are critical for the temporal coordination necessary for effective integration of multimodal information (89).

#### 5.1. Coherence Analysis

We assessed individual responses to speech envelope and visual motion by computing magnitude-squared coherence for each trial and electrode using the *mscohere* function in Matlab, applying Welch’s averaged modified periodogram method. The analysis spanned a frequency range from 0.1 to 30 Hz, in increments of 0.33 Hz (90).

The analysis targeted delta (δ, ∼4 Hz) and theta (θ, 4-8 Hz) frequency bands, identifying frequencies where coherence peaked most prominently for each stimulus condition. Statistical comparisons across groups and stimuli were conducted using the clusters identified through the method outlined in Section 4, combined with a nonparametric test and Dunn’s multiple comparisons test (87). Significance of these findings was established using a surrogate-corrected coherence approach. Surrogate distributions were generated by randomly shifting the neural time course relative to the stimulus feature time courses, preserving their original temporal structure. This process was repeated 50 times for each stimulus condition to generate a robust surrogate distribution. The resulting coherence values were then standardized (Z-scored) against this distribution.

#### 5.2. Phase Analysis

We also performed a phase analysis by calculating the cross-power spectral density (CPSD) phase for each stimulus, electrode, and trial. This was done using the *cpsd* function in Matlab, employing parameters consistent with the coherence analyses. Phase values were determined based on the peak frequency identified in the coherence analysis. Group comparisons were conducted using Matlab’s Circular Statistics Toolbox (91). A two-way parametric ANOVA for circular data was performed to facilitate a nuanced comparison between pairs of conditions and groups, followed by post-hoc comparisons using the Watson-Williams multi-sample test (91). Meanwhile, the Rayleigh test was performed to investigate whether phase distribution is under unimodal distribution. Our focus was primarily on electrodes identified through TRF estimation outcomes.

## Results

### 1.1 Atypical neural tracking of speech envelope in ASD children

We found distinct neural tracking for the auditory and visual parts of the stimuli. Different scalp distribution patterns were observed between the two groups (Figure 2). The ASD group had reduced neural response to the speech envelope relative to the TD group (Figure 2A, top row). In contrast, there was no difference in visual motion processing between the two groups (Figure 2A middle row). These results suggest atypical auditory but not visual processing of dynamic communicative stimuli in young children diagnosed with ASD.

### 1.2 Univariate decoding models confirm atypical speech envelope processing in children with ASD

Stimulus reconstruction accuracy was different in the ASD and TD groups. While reconstruction accuracy was equivalent for speech envelope and visual motion (p>0.9999, Figure 2B) in the TD group, it was lower for speech than for visual motion (p<0.0001, Figure 2B) in the ASD group. Group comparison shows that speech envelope reconstruction was weaker in the ASD than in the TD group (p <0.0001, Figure 2B), without significant group difference for visual motion reconstruction (Figure 2B). These results align with neural tracking results to suggest that it is mostly speech processing that is impaired in autistic children.

### 2. Multivariate encoding models indicate atypical audiovisual processing in ASD children

Further, we explored whether speech anomalies in ASD are limited to auditory processing difficulties or associated with audiovisual (AV) processing anomalies. We found stronger neural representation of the combined AV stimulus compared to individual single stimuli in both groups, an expected result as multivariate models provide in general more accurate prediction of neural responses by combining information from multiple sources. More interestingly, the joint model suggested weaker AV representation in the ASD than in the TD group (Figure 2A bottom row).

### 3. Compromised audiovisual integration in ASD disrupts visual enhancement of auditory processing

By comparing the decoding accuracies for speech envelope and visual motion across univariate (A-only and V-only) and multivariate (AV-joint) models, we found distinct patterns of audiovisual integration in ASD and TD children (Table 1, Figure 3). Notably, in the TD group, the accuracy of speech envelope reconstruction in the AV-joint model (concurrent speech envelope and visual motion) did not significantly differ from the A-only model (p = 0.1414, Figure 3A). Conversely, in the ASD group, the speech envelope was decoded with significantly less accuracy in the AV-joint model than in the A-only model (p=0.0304, Figure 3B), indicating a disruptive effect of AV integration on auditory processing specific to this group. Despite these differences, both groups exhibited decreased visual motion reconstruction accuracy in the AV-joint model compared to the V-only model (ASD group: p < 0.0001, TD group: p = 0.0373, Figure 3), with a more pronounced decrement observed in the ASD group (p = 0.0003, Table 2). These findings suggest that while AV integration generally impacts visual processing across both groups, it adversely affects auditory processing primarily in the ASD group.

**Figure 3.**
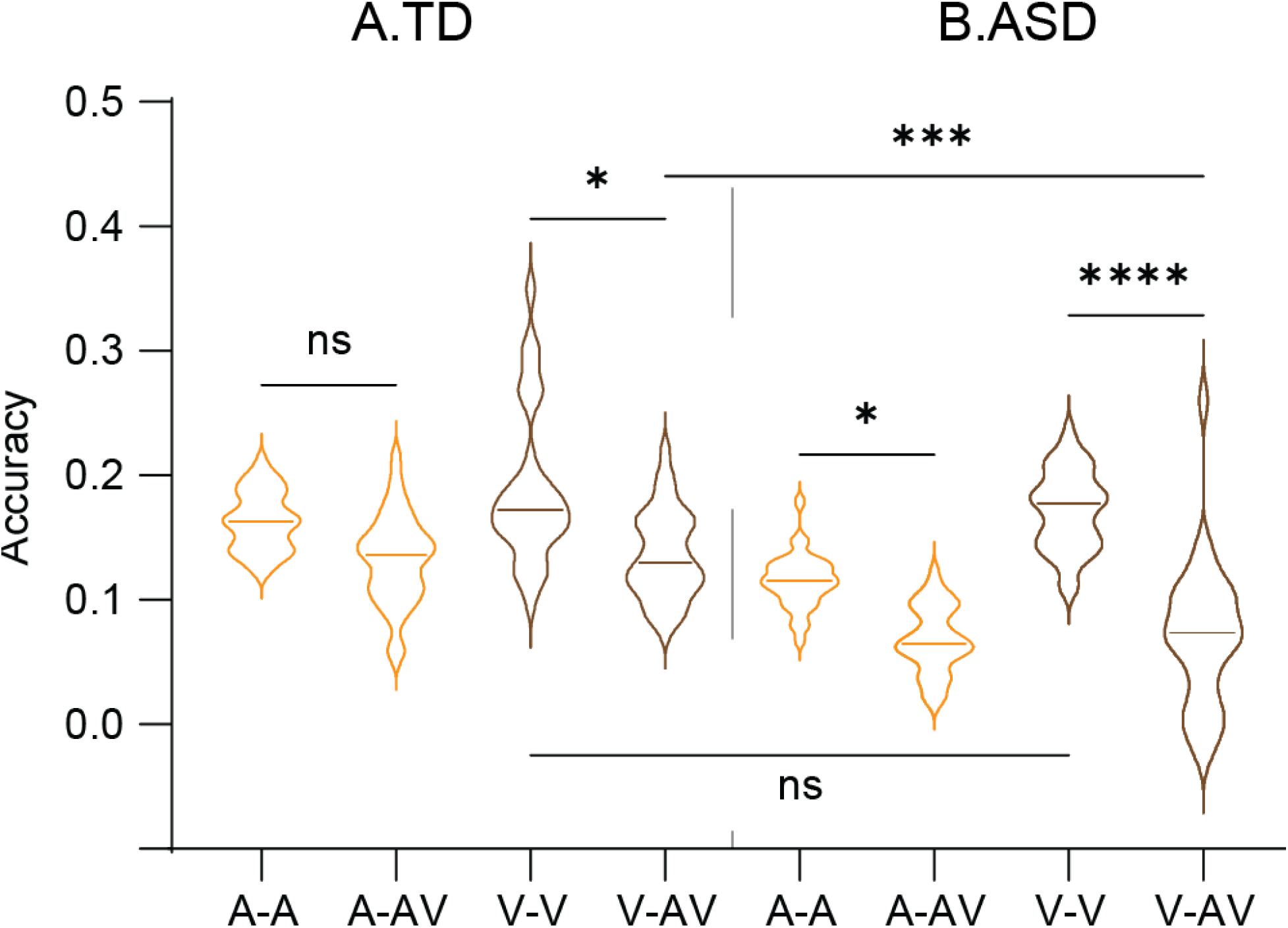
Evaluation of decoding accuracy in ASD (A) and TD (B). Stimulus reconstruction accuracy: speech envelope(A-) and visual motion(V-) in both the single-stimulus model (A-only = A-A and V-only = V-V) and the AV-joint model (A-AV, and V-AV)). Significance levels are indicated as follows: ‘ns’ for p>0.05 (not significant), * for p <0.05, ** for p<0.01, *** for p<0.001, **** for p<0.0001. For additional details, see Supplemental Figure 3*-1*.

**Table 1.**
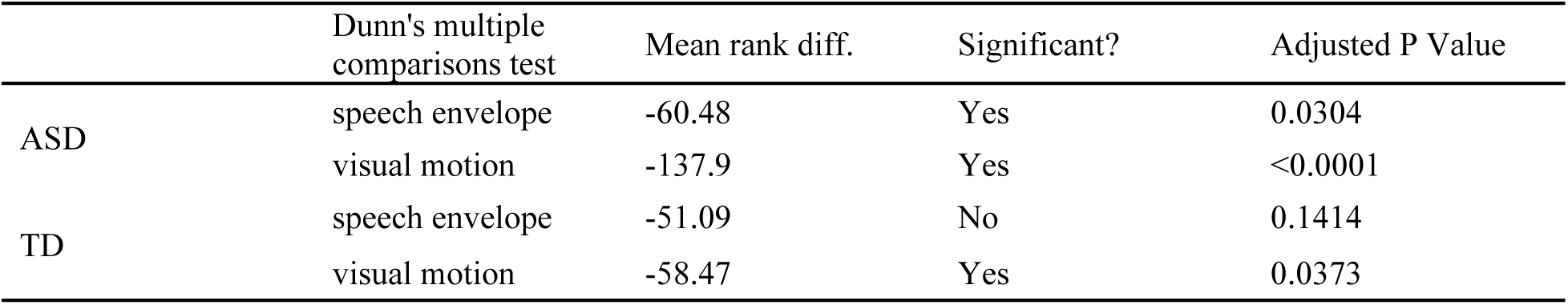
The statistical difference among joint model and single models for speech envelope and visual motion.

|  | Dunn's multiple comparisons test | Mean rank diff. | Significant? | Adjusted P Value |
| --- | --- | --- | --- | --- |
| ASD | speech envelope | -60.48 | Yes | 0.0304 |
|  | visual motion | -137.9 | Yes | <0.0001 |
| TD | speech envelope | -51.09 | No | 0.1414 |
|  | visual motion | -58.47 | Yes | 0.0373 |

**Table 2.** The statistical difference of the decoding accuracy evolution for speech envelope and visual motion.

|  | Dunn's multiple comparisons test | Mean rank diff. | Significant? | Adjusted P Value |
| --- | --- | --- | --- | --- |
| Visual motion | ASD v.s.TD | -37.55 | Yes | 0.0003 |
| Speech envelope | ASD v.s.TD | -13.32 | No | 0.8866 |
| TD | V v.s. A | -6.094 | No | >0.9999 |
| ASD | V v.s. A | -30.32 | Yes | 0.0065 |

A: speech envelope
V: visual motion

### 4. No visual precedence in audiovisual processing in ASD

When studying the temporal dynamics of auditory and visual processing to assess audiovisual integration, we found distinct time-lags in auditory and visual decoding accuracies in ASD and TD children. In the TD group, it took 200∼ms to reach significant decoding accuracy for auditory responses but only ∼50ms for visual ones. The exact opposite pattern was found in the ASD group with a ∼200 ms time-lag got visual decoding versus ∼50 ms for auditory decoding (Figure 4A). This analysis revealed a visual lead in the TD consistent with the fact that speech sources are usually ahead of sounds, but an auditory lead in the ASD group (Figure 4B). This temporal inversion indicates a fundamental alteration in the sensory processing sequence for children with ASD.

**Figure 4.**
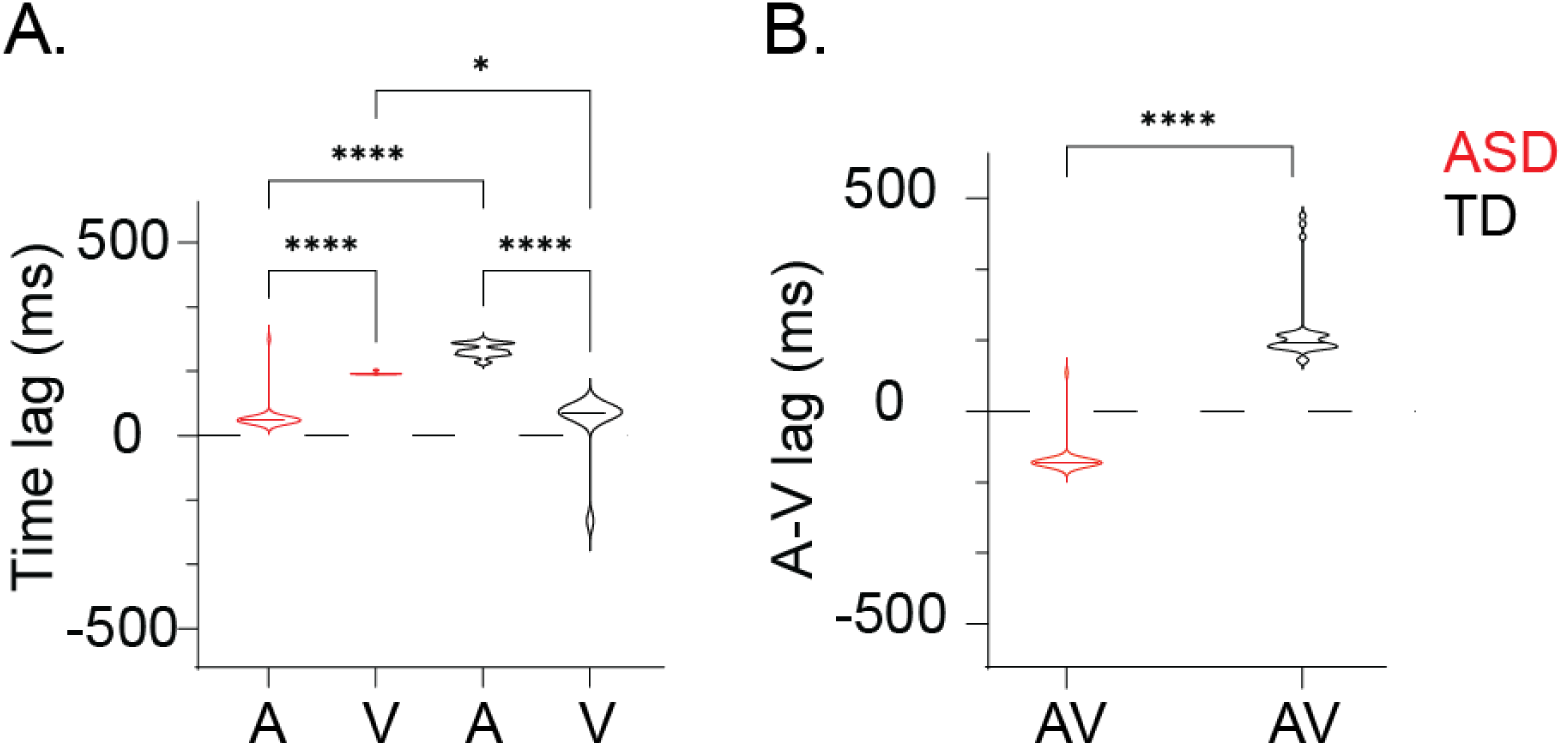
Optimal EEG-stimuli time lag for ASD (red) and TD (black) groups. (A) depicts the optimal time-lag observed in the reconstruction of stimulus features in AV-joint model, specifically speech envelope (A-) and visual motion(V-); Positive values represent stimulus lead EEG signal. (B) illustrates the A-V time lag AV-joint model. Positive values represent V leads A. Significance levels are indicated as follows: ns>0.05, *p <0.05, **p<0.01, ***p<0.001, ****p<0.0001.

### 5. Intact capacity to synchronize neural activity to the stimulus input in ASD

To ascertain whether speech tracking anomalies in ASD are primarily attributable to a general defect in stimulus/brain synchronization or rather results from audiovisual integration deficits, we investigated the intricate temporal dynamics underlying these processes. Specifically, we explored the coherence of stimulus-response relationships within delta (1–4 Hz) and theta (4–8 Hz) frequency bands, typically associated with syllable-and phrase-level speech processing. We observed a higher stimulus/brain coherence in the delta band than the theta band, yet when comparing across groups, both the average coherence values and their distribution patterns showed remarkable similarity (Figure 5, Table 3). This uniformity shows that the intrinsic capacity of neural activity to synchronize with auditory and visual stimuli is consistent between groups (ASD v.s. TD).

**Figure 5.**
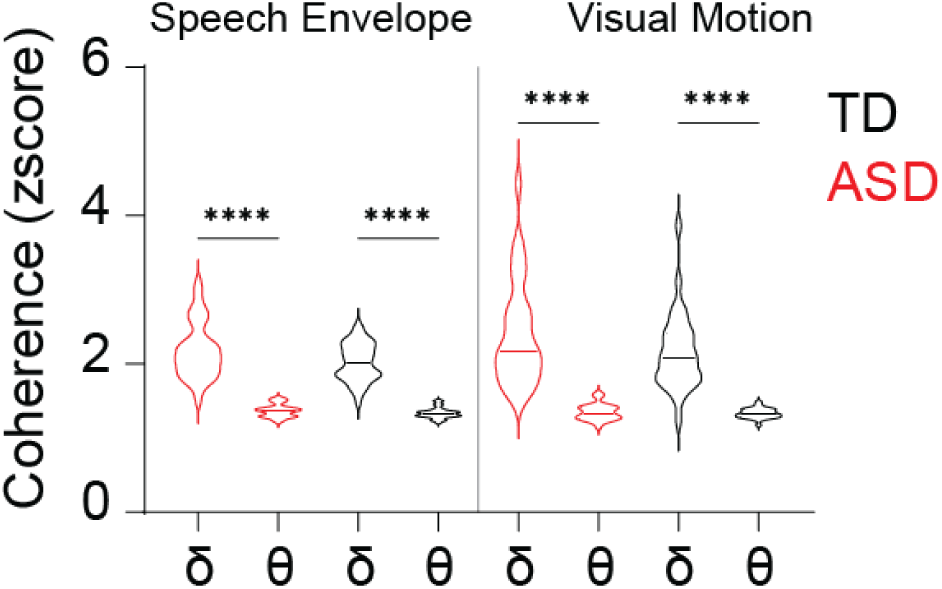
**Stimulus-response coherence** in Theta and Delta Bands for ASD (red) and TD (black) groups. The plot displays the coherence between stimulus and response for Speech Envelope and Visual Motion. Error bars represent the standard error of the mean. The coherence levels are compared within the specific frequency bands of interest, highlighting potential group differences in sensory processing. Significance levels are indicated as follows: ns>0.05, *p <0.05, **p<0.01, ***p<0.001, ****p<0.0001.

**Table 3.** The statistical difference across groups and frequency bands in stimulus-response coherence.

|  |  | Dunn's multiple comparisons test | Mean rank diff. | Significant? | Adjusted P Value |
| --- | --- | --- | --- | --- | --- |
| ASD | A | delta v.s. theta | 162.7 | Yes | <0.0001 |
| ASD | V | delta v.s. theta | 191.5 | Yes | <0.0001 |
| ASD | delta | A v.s. V | -7.774 | No | >0.9999 |
| ASD | theta | A v.s. V | 20.94 | No | >0.9999 |
| TD | A | delta v.s. theta | 178.6 | Yes | <0.0001 |
| TD | V | delta v.s. theta | 176.4 | Yes | <0.0001 |
| TD | delta | A v.s. V | -0.6563 | No | >0.9999 |
| TD | theta | A v.s. V | -2.844 | No | >0.9999 |
| A | delta | ASD v.s. TD | 12.3 | No | >0.9999 |
| A | theta | ASD v.s. TD | 28.19 | No | >0.9999 |
| V | delta | ASD v.s. TD | 19.42 | No | >0.9999 |
| V | theta | ASD v.s. TD | 4.406 | No | >0.9999 |

A: speech envelope
V: visual motion

### 6. Theta-range desynchronization of audio-visual responses in ASD

Given the preserved synchronization capacity between brain activity and external stimuli in each modality, we then sought whether audio-visual integration anomalies, notably the inverted AV temporal patterns, are associated with a phase desynchronization of auditory and visual processing. In the delta band, we observed similar phase angles for both groups (F(1,62) = 0.494, p < 0.470), indicating comparable phase locking at this slower frequency, suggesting that temporal alignment in this low-frequency range does not differentiate between groups. Moreover, the small angle indicates the absence of delta band phase-shift between modalities. Yet, a significant group difference was observed in the theta band. The TD group showed a phase-shift of approximately 90 degrees, signaling effective sequential integration with one modality leading the other by a consistent temporal offset, that optimizes audio-visual integration at the syllable level. In ASD children the phase-shift amounted to 180 degrees and the group difference was significant (F(1,62) = 12.05, p < 0.001) (Figure 6). The observed 180-degree phase shift in ASD could suggest that auditory and visual information is out of sync: when one sensory modality is at its peak processing efficiency, the other is at its lowest, potentially leading to disjointed even conflicting sensory processes.

**Figure 6.**
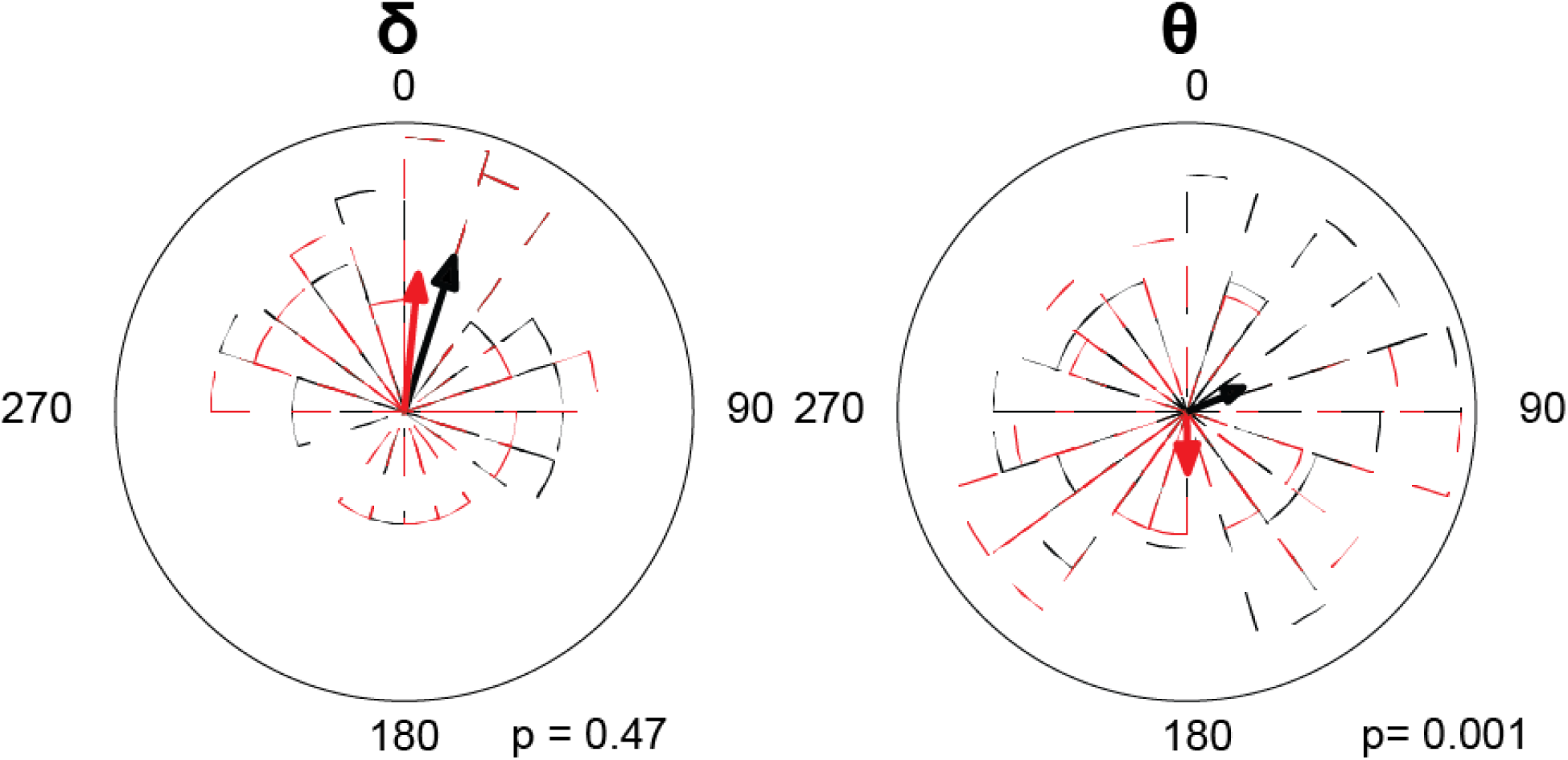
Phase-shift distribution between speech envelope and visual motion. This figure shows the phase-shift distribution between the brain processes of speech envelope and visual motion stimuli for each group. The circular mean of the phase-shift across all subjects is indicated by colored lines: red for the ASD group and black for the TD group. Corresponding polar histograms in red (ASD) and black (TD) visually represent the distribution of phase-shifts for each group. Both groups were tested against the hypothetical uniform distribution of delta (rayleigh test, ASD: p <0.001, rayleigh r = 0.98, TD: p <0.001, rayleigh r = 0.98) and theta phase (rayleigh test, ASD: p <0.001, rayleigh r = 0.95, TD: p <0.001, rayleigh r = 0.96).

### 7. AV phase-shift is related to auditory encoding accuracy in TD and visual encoding accuracy in ASD

Finally, we sought to understand the relationship between the theta band AV phase-shift and the accuracy of auditory and visual information reconstruction within an unimodal framework (Figure 7). As could be expected, in TD children the AV phase-shift did not influence visual reconstruction accuracy (r = 0.033, p = 0.858), but there was a weak negative correlation between the phase-shift extent and speech reconstruction accuracy (r = −0.272, p = 0.132): when the AV phase-shift increased speech reconstruction accuracy decreased, which given the visual lead previously observed could suggest a causal effect. A different pattern was seen in ASD children, with no relation between the phase shift extent and speech reconstruction accuracy (r = 0.197, p = 0.288), but a weak negative correlation between the phase-shift extent and the accuracy of visual information reconstruction (r = −0.325, p = 0.074), with larger AV phase-shifts linked to poorer visual reconstruction accuracy. Likewise, given the auditory lead observed in children with ASD, this could suggest a causal effect.

**Figure 7.**
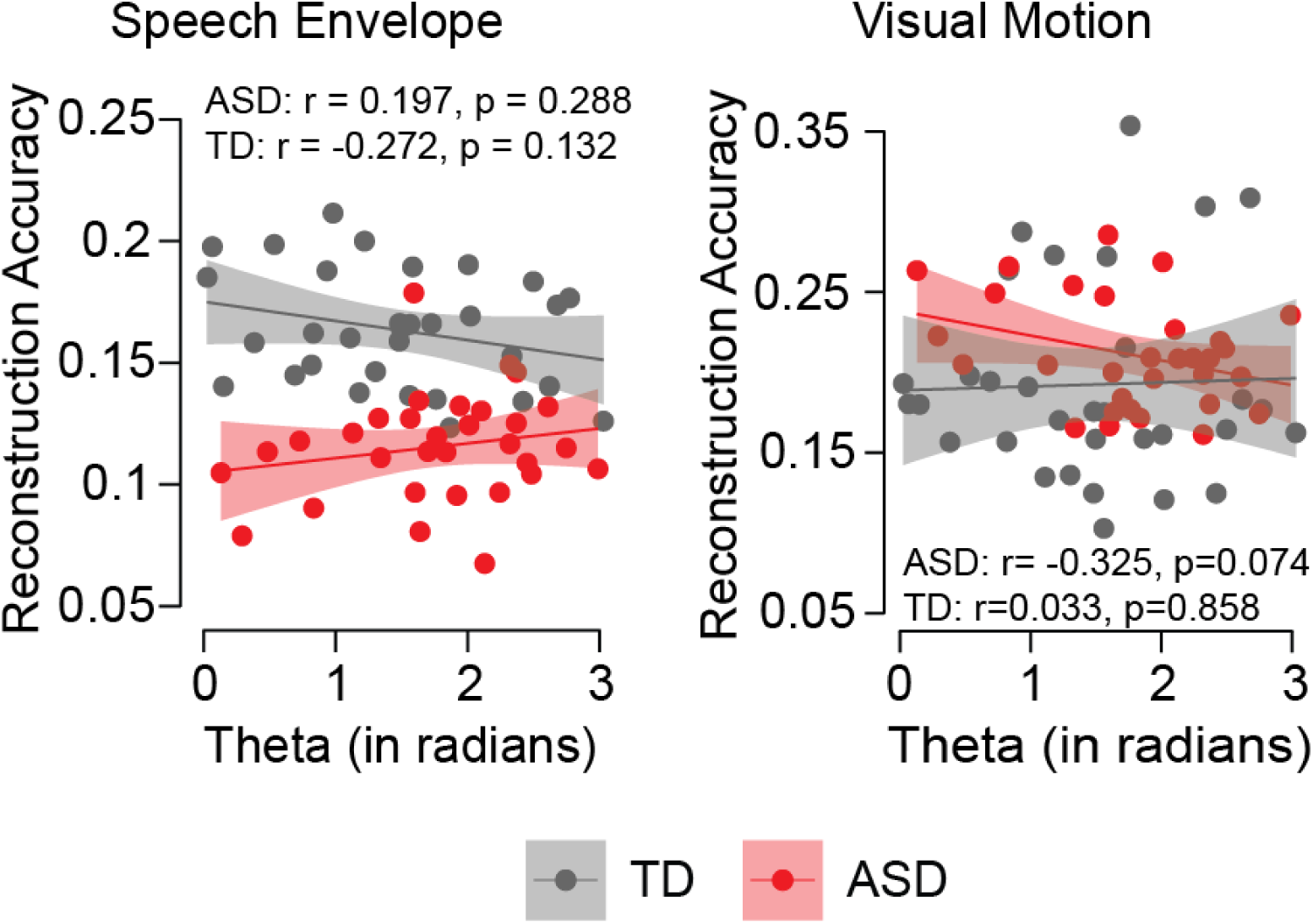
**The relationship between theta phase-shift and reconstruction accuracy** of speech envelope (left) and visual motion (right) in ASD and TD. ASD group suggests a greater phase shift between speech envelope and visual motion positively correlates with speech reconstruction accuracy but negatively correlates with visual reconstruction, while reversely in TD group.

Contrasting patterns in TD and ASD children underscore distinct audiovisual integration mechanisms. In logic, reconstruction accuracy is a proxy of sensory encoding accuracy. Thus, in TD children, the data confirm the known reliance on visual cues to enhance auditory processing, with any misalignments adversely affecting speech information integration. Conversely, in ASD children, while speech encoding is weaker and overall less dependent on visual-auditory phase congruency, visual processing is vulnerable to strong AV desynchronization.

## Discussion

Using several analyses of the EEG recorded in very young children with and without ASD while they were watching a short animated movie, we confirmed previous results showing profound anomalies of the capacity to follow speech rhythms (24,92), an essential prerequisite to speech comprehension. The present study goes beyond this observation by showing that children with ASD did not exhibit the natural dominance of auditory processing when exposed to natural audio-visual speech conditions. Instead, their processing of audio-visual stimuli was impacted by a temporal misalignment of these sensory inputs, which disturbs the predictive processing typically at play when perceiving speech.

### Audio-visual integration anomalies interfere with sensory encoding in ASD

Audio-visual processing, especially the synchronization of the two sensory modalities, plays a pivotal role in understanding the communicative challenges observed in ASD. Previous studies have established that basic auditory dysfunctions and atypical speech processing are characteristic of ASD from an early developmental stage (19–24). Our study reveals that such anomalies are not merely isolated auditory deficits but are deeply connected to the integration of auditory and visual information, a process critical for effective communication, particularly in dynamic or complex listening environments (93,94).

Our findings reveal a specific disruption in audio-visual integration among children with ASD, manifesting in visual dominance and temporal disorganization in auditory and visual processing. This disruption sharply contrasts with the expected auditory processing dominance (95) and might significantly contribute to the language development anomalies encountered by these children. In typical development, the precedence of orofacial visual cues during speech facilitates auditory comprehension through predictive processing, optimizing the brain’s synchronization to incoming speech signals (96). In ASD, extended integration time windows and the lack of effective synchronization of auditory responses by visual signals (as evidenced by the atypical theta band phase-shifts) suggest they cannot use visual cues to facilitate auditory speech processing. On the contrary, our findings show that auditory cues perturb visual processing of communicative situations.

### Specific repercussions of disrupted audio-visual processing on speech tracking

Children with ASD demonstrate effective visual motion tracking and processing capabilities, comparable to their TD peers, despite distinctive scene analysis patterns as previously observed (13,97). Within their preferred exploration zones, children with ASD process visual motion similarly to their TD peers (97), exhibiting a level of bottom-up excitability to visual stimuli akin to TD children (98,99). Our univariate encoding results suggest that the neural activity responsible for visual motion tracking operates similarly in both ASD and TD groups.

However, when visual processing co occurs with speech processing, some difficulties appear. Our multivariate modeling indicates that the neural encoding of audio-visual percepts in ASD children is less efficient, confirming that audiovisual contexts can disrupt brain responses to speech in this population (100). In the same vein, Shic et al. (2020) underscore such AV integration difficulties, noting that children with ASD tend to attend less to faces and mouths in general and more specifically when they produce speech (7). Our study reinforces this crucial finding by showing that while children with ASD encode single visual streams relatively well (visual motion tracking in univariate modeling), they struggle with the concurrent encoding of both auditory and visual streams (multivariate modeling).

Building upon this framework, the research conducted by Chawarska et al. (2022) reveals that 12-month-old infants at risk for ASD, even though they explore faces and mouths similarly to infants with no family history of autism (101), cannot leverage audiovisual cues for language acquisition as do typical children. Our study uncovers the potential underpinnings of the audiovisual integration difficulties observed in ASD. The decoding results indicate that while audio-visual integration interferes with visual processing in both ASD and TD groups, its impact on speech processing is particularly detrimental in the ASD group. Thus, the impairments in AV integration we observe are not merely additive but they interactively exacerbate sensory processing challenges much more adversely in children with ASD.

### Audio-visual temporal integration underlies speech impairment in ASD

Audiovisual integration capitalizes on the temporal alignment of sensory events, with visual information often enhancing the auditory signal’s clarity and precision, especially when auditory cues are poor, noisy, or ambiguous (102–104). Visual cues related to speech are typically processed faster than auditory cues, allowing visual information to facilitate synchronizing subsequent auditory processing (105). Our findings confirm in TD children a visual lead (∼50 ms) within a temporal window is conducive to effective interaction and coordination between auditory and visual cues. This window reasonably aligns with established models, positing a 200 ms integration period (39,61–63), ranging from a 30 ms visual lag to a 170 ms of visual lead (61).

The precise timing of audio-visual sequences is fundamental to audio-visual integration via predictive processing, whereby the brain leverages visual cues to anticipate and decode forthcoming auditory information. Here, phase-locking analyses in TD children show that the neural responses associated with auditory and visual processing exhibit a 90-degree phase shift. This observation indicates that the brain orchestrates visual and auditory information in a synergistic but temporally distinct manner. Such a phase relationship is instrumental in achieving a dynamic balance between the sensory streams, facilitating an integration that enhances perception and communication (106,107). The pivotal role of the theta frequency band in orchestrating audio-visual speech processing is robustly supported in the literature (56,58,59). A pi/2 phase shift during an audio-visual speech event might fine-tune the phase alignment to a timing that is congruent with the auditory inputs. This meticulous phase alignment contrasts sharply with the broad phase distribution observed in the ASD group, hinting at a pronounced disparity in how auditory and visual cues are integrated.

Crucially, in ASD children, the integration of auditory and visual streams is jeopardized, as evidenced by our observation of an atypical auditory lead (∼50 ms), which disrupts the conventional sequence where visual information typically precedes auditory. This inversion undermines the usual enhancement provided by visual signals to auditory processing, highlighting a marked alteration or impairment in multisensory integration. Furthermore, we noted a 180-degree phase-shift in the neural activities associated with processing these 2 streams. This substantial phase misalignment reflects a profound disruption in temporal coordination, potentially leading to confusion or interpretation errors. Such a discrepancy underscores a critical deficiency in predictive processing in ASD, where, rather than synergistically enhancing each other, auditory and visual cues conflict, undermining the synthesis of coherent audio-visual perception. This misalignment is also reflected in the broader phase distribution seen in children with ASD, suggesting that they might require an extended temporal window to reach effective processing (31,32).

Our results thus confirm that the phase of low-frequency neural oscillations is crucial for the encoding of order - for instance with the implication of the theta band in working memory (108) or for temporal parsing in speech (109). The anomaly in temporal encoding mechanisms described in our experiment is constrained by the temporal features provided by external stimulation to build a temporal reference frame. While delta oscillations have previously been linked to temporal predictability (110,111), we observed here that sensory integration is affected by AV misalignment in the theta range, which is associated with atypical speech perception in ASD. The AV integration primarily occurs at the syllable level with a typical tolerance of AV asynchrony at 250ms, which corresponds to the theta range (39,61–64).

## Conclusion

We show remarkable anomalies in audio-visual integration in children with ASD. We confirm previous findings of disrupted speech rhythm tracking and further reveal a specific disruption in audio-visual integration, manifesting as temporal desynchronization. This disruption significantly impacts speech processing, contributing to the communicative challenges faced by children with ASD. Our results highlight the critical role of temporal processing in audio-visual integration and underscore the importance of characterizing these mechanisms in ASD. Moving forward, these insights could inform the development of targeted interventions aiming at regulating temporal speech processing and AV synchronization to improve communication in children with ASD.

## Author Contributions

Data acquisition and clinical resources: M.S., N. K.; Study design: X.W., N. K., M.S. and A-L.G.; Data analysis: X.W., S.B.; Writing-original draft: X.W.; Writing-review-editing: X.W., S.B., N. K., A-L.G. and M.S.; Supervision: A-L.G., and M.S.

## Data and materials availability

The unprocessed datasets for this manuscript are not publicly available yet, because analysis of these data is ongoing as part of a longitudinal study, and results are expected to be published in the future. When all data has been published, requests to access the datasets should be directed to Dr Marie Schaer, the custom MATLAB analysis scripts will be made available upon request to the Lead Contact, Xiaoyue Wang.

