## Supplemental Figure 3 for "Atypical Audio-Visual Neural Synchrony and Speech Processing in children with Autism Spectrum Disorder"

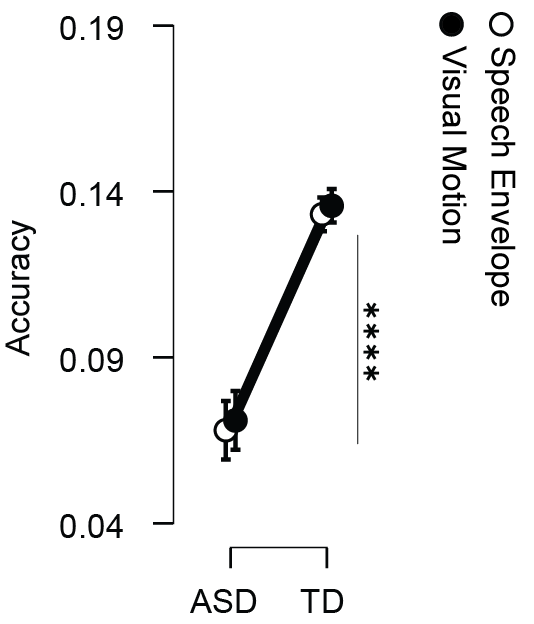

*Supplement Figure 3-1 ASD and TD groups' decoding accuracy in the AV-joint model. This figure presents the reconstruction accuracies for stimuli, specifically the speech envelope and visual motion from AV joint model. The error bars represent the standard error of the mean. Significance levels are indicated as follows:‘ns’ for p>0.05 (not significant), * for p <0.05, ** for p<0.01, *** for p<0.001, **** for p<0.0001.*
